# Autophagy in DG engrams mediates Rac1-dependent forgetting via TLR2/4 signals in microglia

**DOI:** 10.1101/2021.08.26.457763

**Authors:** Zhilin Wang, Ruyan Chen, Qing Lin, Yan Jiang, Qiumin Le, Xing Liu, Lan Ma, Feifei Wang

## Abstract

**BACKGROUND:** Engrams are considered to be substrates for memory storage, and the adaptive plasticity remolding that switch engrams to an inaccessible state cause forgetting. The normal function of engrams is ensured by the crosstalk between neurons and microglia, the major immune cells in the brain implicated in synapse remodeling and memory processing. However, the cellular processes and molecular mediators between engrams and microglia underlying forgetting are poorly understood.

**METHODS:** We utilized doxycycline (Dox)-dependent robust activity marking (RAM) system to label and manipulate DG engrams encoding contexture fear memory in mice. Combining optogenetics, microglia-specific transcriptomics, fluorescence in situ hybridization and spine morphology analysis, we investigated the potential mechanisms of information exchange between engrams and microglia mediating memory forgetting.

**RESULTS:** The expression of *Rac1* in dentate gyrus (DG) engrams upregulated memory encoding. Increased Rac1 activity in DG engrams accelerated forgetting, upregulated autophagy influx and the expression of autophagy protein 7 (*Atg7*). The elevated ATG7 expression in the engrams activated of DG microglia and promoted forgetting. In addition, the Toll-like receptor (TLR) signal in DG microglia was upregulated when overexpressing ATG7 or activating Rac1 in DG engrams, and mediated the ATG7-dependent synapse remodeling and Rac1-dependent forgetting.

**CONCLUSIONS:** In this study, we found that Rac1 increases ATG7-dependent autophagy in DG engrams, which activated microglia via TLR2/4, to promote spine remodeling and forgetting. These results unravel a novel pathway mediating memory forgetting, and provide a potential therapeutic strategy in the treatment of cognitive disorders such as Alzheimer’s disease.

## Introduction

Learning ability helps us encode information and form specific memories to promote adaptive behavior in an ever-changing environment. A given memory is encoded by sparsely distributed engrams which are a population of neurons activated by learning and engaged enduring physical and/or chemical changes in the brain(1). However, when memory is consolidated, forgetting happens at the same time. The recently proposed concept that forgetting is a higher order of learning, enhancing the utility of memory by altering engram cell accessibility, underlies the importance of forgetting to an organism (2). Numerous researches have been proposed to explain the putative mechanisms of forgetting (2), including changes in synaptic plasticity modulated by AMPA receptors trafficking, spines morphology, and the other cells such as microglia and astrocyte. However, causal evidence for interaction between engrams and the surroundings contributing to forgetting is still lacking.

Previous researches have shown that microglia, the major immune cells in the brain, are involved in both neurogenesis-related and aging-dependent memory destabilization(3–5) and microglia-neuron interactions play a role in regulating forgetting via complement-and activity-dependent synaptic elimination (3). Microglia become morphologically and functionally activated by other brain-derived materials, such as protein deposits, and axonal and synaptic structures. Toll-like receptors (TLRs), pattern recognition receptors for pathogen-associated molecular patterns (PAMPS), mediate activation of microglia and play an important role in inflammation and immune responses in the CNS (6, 7). TLRs-mediated signaling pathways in microglia lead to the translocation of transcription factors, such as NF-κB, thereby activating the transcription of several genes including interleukins (ILs) and tumor necrosis factors (TNFs) which mediate synaptic pruning and functional plasticity in neurons (8–10). Nevertheless, the relationship between activation of microglia via TLRs and the memory forgetting caused by improper synapse remodeling has not been investigated. This is crucial for figuring out the possible pathogenesis of psychiatric disorders because of the impaired synaptic refinement, such as Alzheimer’s disease (11, 12), schizophrenia (13, 14) and autism (15).

It has been reported that retrieval of a consolidated memory can destabilize the memory, after which gene transcription and new protein synthesis are required for memory stabilization and reconsolidation (16–18). That indicates protein degradation, the flip of protein synthesis, may also plays a role in memory reorganization because it can remove existing proteins and incorporate new proteins into the synapses. Indeed, it has been reported that inhibition of the ubiquitin- and proteasome-dependent protein degradation prevents memory destabilization (19), while induction of synaptic protein degradation through autophagy, another protein degradation pathway, can enhance memory destabilization upon retrieval (20). Moreover, autophagy is regulated by learning and neuronal activity and contributes to directing internalized AMPAR degradation, which may weaken synaptic connectivity among engrams, leading to less successful reactivation and forgetting (20–22). These observations got us interested in investigating whether autophagy in engrams would affect memory reorganization and contribute to forgetting. Autophagy is a complex self-degradative system that removes damaged cellular debris and aggregated proteins to maintain brain homeostasis and promote neuronal survival (23). In addition to the conventional homeostatic and adaptive function, autophagy is also necessary for immune function by regulating cytokine production and release, inflammasome activation, and clearance of invading pathogens. Exactly, recent researches have shown that autophagy is involved in microglia activation in an unconventional secretion pathway (24). On the basis of these studies, we launched an effort to investigate whether autophagy in engrams could activate microglia and then acted as a node mediating forgetting induced by microglial remodeling of synapses.

In this study, we found that the *Rac1* mRNA and autophagy protein 7 (ATG7) in the dorsal dentate gyrus (DG) engrams were both increased after contexture fear consolidated, and the elevated signaling both led to the impairment of fear memory retrieval through TLR2/4-dependent activation of microglia. In addition, increased Rac1 activity also promoted autophagy induction in DG engrams. These interactions among Rac1, ATG7 and crosstalk between engrams and microglia lead to synapse remodeling of engrams and memory destabilization, which might help us get insight into the mechanism of forgetting and be potential to be used as a therapeutic tool in the treatment of cognitive impairment.

## Results

### Forgetting signal Rac1 in DG engrams impairs fear memory and promoted autophagy and the expression of ATG7

Elevated Rac1 activity in the hippocampus has been reported to led to the forgetting of different memories(25–28). To investigate the expression and effects of Rac1 in DG engram in contexture fear memory, a doxycycline (Dox)-dependent robust activity marking (RAM) system was used to label the engrams (29). The specific function of the engrams labeled by the RAM system was verified by expressing the inhibitory (hM4Di) or excitatory (hM3Dq) DREADDs (designer receptors exclusively activated by designer drugs)(30) in the engrams. Clozapine N-oxide (CNO) was intraperitoneally (i.p.) delivered 30 min before the contexture memory tests (Supplemental Figure S1A). The freezing level of mice was bidirectionally regulated by inhibition or activation DG engrams (Supplemental Figure S1B-C). No differences were detected in the open field test (Supplemental Figure S1D), suggesting locomotor activity in mice was not affected by chemogenetic manipulation of DG engrams. The RAM reporter system also labeled small populations of vGluT2 positive cells in the dentate hilus regions (Supplemental Figure S1E). Although these excitatory neuron cells can make synaptic connections with granule cells in DG and integrate numerous inputs from other hippocampal regions, researches have shown that the number of labeled hilar cells was comparable in the home cage and conditioning context (31).

The level of *Rac1* mRNA in dentate gyrus (DG) engrams were assessed by single molecule fluorescence in situ hybridization (sm FISH). We analyzed the *Rac1* mRNA level within the engrams at different time points (4h, 12h, day 1, and day 3) after fear conditioning (Fig 1A-C). Rac1 mRNA expression within DG engrams is upregulated after contextual fear learning. To investigate whether Rac1 activity in DG engrams impairs the contextual fear memory, WT mice were infected with *AAV-RAM -Cre* mixed with either *AAV-DIO -C450A–mVenus*, *AAV-DIO-Pa Rac1-mVenus* or *AAV-DIO-T17N-mVenus* in bilateral dorsal DG to express light-insensitive mutant (C450A), photoactivatable Rac1 (Pa Rac1), or photo inhibitable Rac1 (T17N) in the engrams (Supplemental Figure S1F). 3 days after engrams labeling, the light pulse (150 ms; 1 Hz; 1 hr) was delivered into dorsal DG 1hr before the retrieval of contextual fear memory (Fig 1D). We noted that the contextual fear memory was significantly impaired after optical activation of Pa-Rac1 in DG engrams, and this effect lasted up to 24 hrs (Fig 1E-F). Instead, optical inhibition of Rac1 T17N within the engrams increased the freezing level in the fear context, but the enhancement was not observed 24 hrs after Rac1 inhibition (Fig 1G-H). Thus, the Rac1 activation accelerated forgetting of contextual fear memory and this result was reversible through inhibition of Rac1 activity. Moreover, Rac1 induced effect on memory maintenance was long-term up to 24h. Additionally, c-Fos immunostaining was performed 1.5 hrs after the memory retrieval. The percentage of the c-Fos^+^ component was decreased in the engrams overexpressing Pa-Rac1 in comparison with the C450A group, suggesting optical activation of Rac1 in DG engrams decreased the reactivation of engrams during memory retrieval (Fig 1I-J).

**Figure 1.**
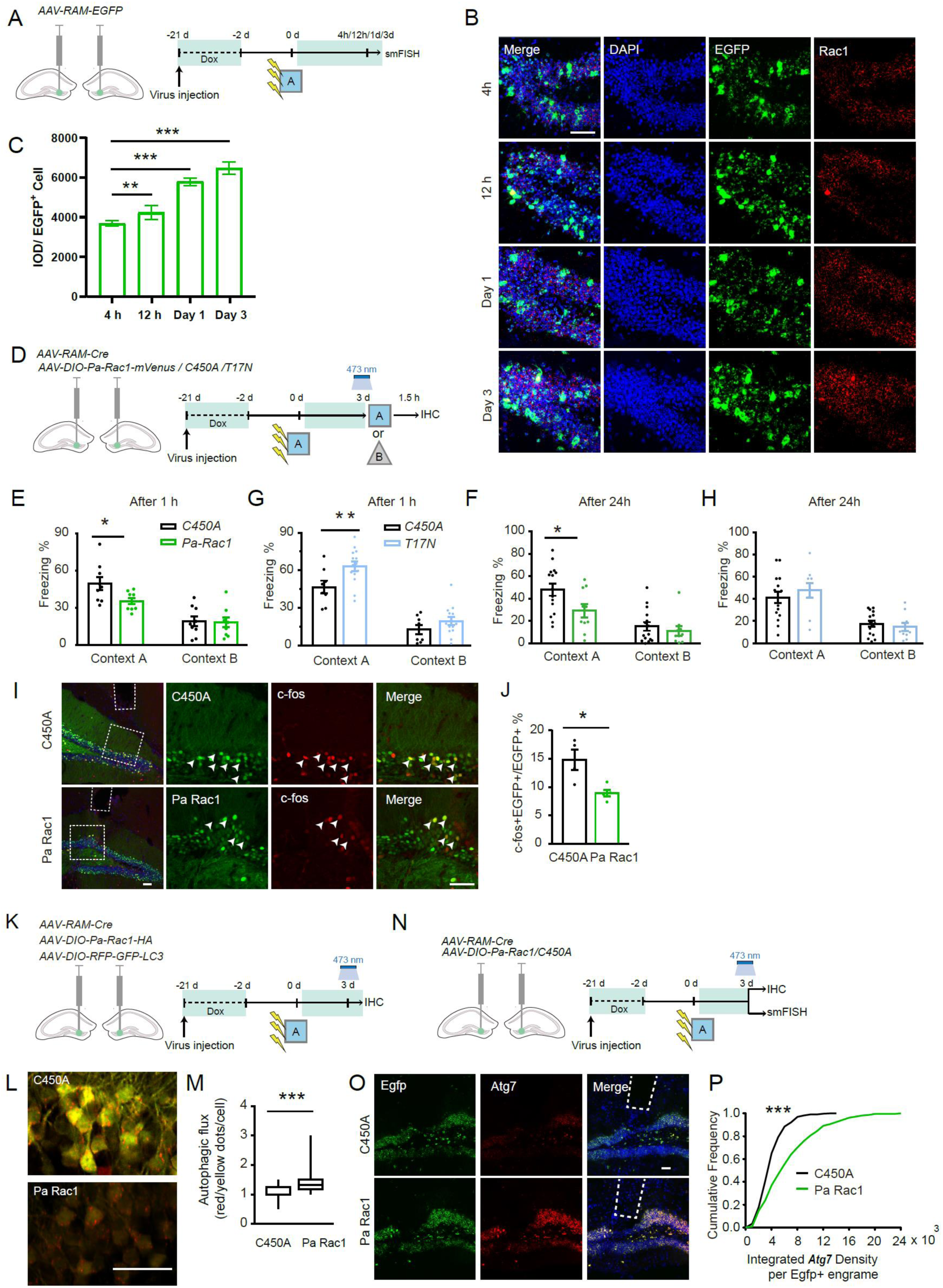
Rac1 in DG engrams is upregulated after memory consolidation, and leads to the impairment of fear memory and enhancement of autophagy of the engrams. **(A)** Experimental procedure to test Rac1 expression after CFC-induced learning. **(B, C)** Representative images (**B**) and quantification **(C)** of the single-molecule fluorescence in situ hybridization (smFISH) intensity of *Rac1* in *Egfp^+^* cell. Kruskal-Wallis H test, χ^2^ =136.056, P<0.001. Green: *Egfp*; Red: *Rac1*; Blue: DAPI. Scale bar:50 μm. **(D)** Experimental procedure to optically activate or inhibit Rac1 within DG engrams. 3 days after engrams labeling, the 473 nm laser pulse was delivered in the DG for 150 ms at 1 Hz for 1 hr, and the freezing level in context A or context B was measured 1 hr or 24h after optical stimulation. **(E)** Freezing level in context A and context B 1 hr after optical stimulation. F_treatment x context (1, 16)_ = 4.531, P=0.049, RM ANOVA. Context A: C450A versus Pa Rac1: P=0.019, Bonferroni post t-test. **(F)** Freezing level in context A and context B 24 hrs after optical stimulation. F_treatment x context (1, 23)_ = 5.06, P=0.034, RM ANOVA. Context A: C450A versus Pa Rac1: P=0.013, Bonferroni post t-test. **(G)** Freezing level in context A and context B 1 hr after optical stimulation. F_treatment x context (1, 20)_ = 2.672, P=0.118, RM ANOVA. Context A: C450A versus T17N: P=0.006. Bonferroni post t test. **(H)** Freezing level in context A and context B 24 hrs after optical stimulation. F_treatment x context (1, 23)_ = 1.116, P=0.302, RM ANOVA. **(I, J)** Brains were collected 1.5 hrs after re-exposure to context A for c-Fos staining. Representative images **(I)** and quantification **(J)** of the percentage of c-Fos^+^ components in EGFP^+^ engrams. t=3.112, P=0.041, Two-tailed unpaired t-test. Green:C450A/Pa Rac1; Red: c-Fos; Blue: DAPI. Scale bar: 50 μm. **(K)** Experimental procedure to assess autophagy after optical activation of Rac1 in the DG engrams. Representative confocal images **(L)** and quantification **(M)** of autophagy influx 3 days after the engrams labeling. Z=-9.287, P<0.001, Mann-Whitney U test. Red: RFP; Green: EGFP. Scale bar: 50 μm. **(N)** Experimental procedure to assess the *Atg7* after optical activation of Rac1 in DG engrams. Representative images **(O)** and quantification **(P)** of the smFish against *Atg7* mRNA in C450A and Pa Rac1 groups. Z=3.872, P<0.001, Kolmogorov–Smirnov test. Green: *Egfp*; Red: *Atg7*; Blue: DAPI. Scale bar: 50 μm. Data are presented as mean ± SEM. *P < 0.05, **P < 0.01, ***P < 0.001.

Previous researches showed that Rac1 induces autophagy through activating JNK signal pathway, which enhances the expression of multiple autophagy-related proteins (ATGs) including Atg5 and Atg7 via a Foxo-dependent transcription pathway(32). To assess whether Rac1 interferes with fear memory via promoting autophagy in DG engram, a Cre-dependent autophagy flux reporter system *AAV-DIO-RFP-GFP-LC3* was conducted(33) based on different pH stability of EGFP and RFP fluorescent proteins (Fig 1K and Supplemental Figure S1G). 1 hr after optical activation of Rac1 within the DG engrams, the autophagy signal was evaluated by analyzing the ratio of RFP and GFP fluorescent intensity. There was a substantial increase on the ratio of the RFP/Yellow puncta in the DG engrams (Fig 1L-M). We further investigated the correlation of Rac1 activation with autophagy induction within engrams by smFISH to assess the ATG7 mRNA level affected by optical activating of Rac1 in the DG engrams (Fig 1N). Notably, optical activation of Rac1 significantly increased the mRNA level of *Atg7* in the DG engrams compared with the C450A control group (Fig 1O-P). Together, these data suggest that upregulation of Rac1 signaling in DG engram accelerates forgetting via facilitating autophagy induction within the engrams.

### ATG7 in DG engrams mediate Rac1-dependent forgetting and microglia activation

Previous researches investigated that autophagy contributes to memory destabilization and enhances the erasure of auditory fear memory (20). Therefore, we were interested in how does autophagy promote memory destabilization? Firstly, we investigated the effect of learning on autophagy because learning-induced stimulations could upregulate hippocampal autophagy (34). The protein level of p62, the substrate of autophagy degraded after fusion with the lysosome, in DG extracts of mice was investigated. The protein level of p62 in the DG was decreased during day 1 to day 3 after fear conditioning (Fig 2A-C). ATG7 is essential for autophagy as it plays a role in autophagosomes fusion with lysosomes. Consistently, the immunostaining of ATG7 showed that the total protein level of ATG7 in DG was increased 1 day after CFC stimulation, and lasted up to day 3 (Fig 2D-E). These data indicated that autophagic flux in DG was increased after memory encoding.

**Figure 2.**
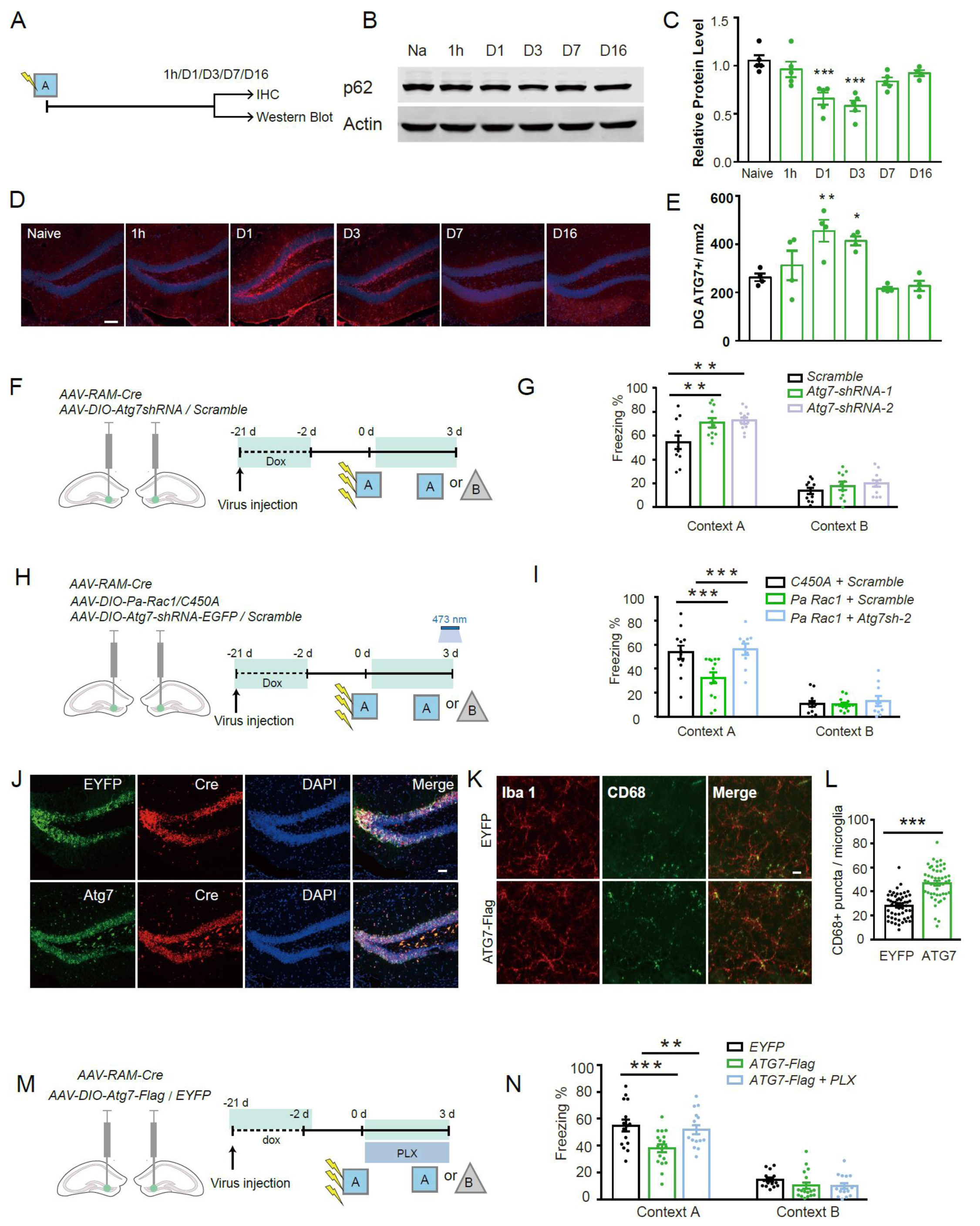
Expression of ATG7 is upregulated after learning and excessive ATG7 in DG engrams leads to the impairment of fear memory and the activation of microglia. **(A-C)** Representative western blot images **(B)** and quantification **(C)** of p62 protein in DG extracts. One-way ANOVA, F=10.084, P<0.001. **(D, E)** Experimental procedure to measure ATG7 protein level in DG. Representative confocal images **(D)** and quantification **(E)** of ATG7 protein in DG. F=18.033, p<0.001, Welch’s test. Red: ATG7; Blue: DAPI. Scale bar: 50 μm. **(F)** Experimental procedure to investigate the effect of ATG7 knockdown in the DG engrams on the contextual fear memory. **(G)** Freezing level 3 days after fear conditioning in context A and neutral context (context B). F_treatment × context (2, 31)_ = 2.330, P=0.114, Repeated measurement ANOVA. Context A: Scramble versus Atg7sh-1. P=0.006; Scramble versus Atg7sh-2, P=0.002; Bonferroni post hoc test. **(H)** Experiment procedure to test the effect of Rac1 activation and ATG7 knockdown within DG engrams on the contextual fear memory. **(I)** Freezing level of mice in context A and context B 3 days after fear conditioning. F_treatment x context (2,33)_ = 6.2, P=0.005, RM ANOVA. Context A: C450A +*Scramble* versus Pa Rac1 + *Scramble*: P<0.001; Pa Rac1 + *Scramble* versus Pa Rac1 + *Atg7sh-2*: P<0.001. Bonferroni post t test. **(J)** Representative confocal images of ATG7 or EYFP expression in Cre^+^ regions. Green: EYFP/ATG7-Flag; Red: Cre; Blue: DAPI. Scale bar:50 μm. **(K, L)** Representative images **(K)** and quantification **(L)** of CD68 puncta within Iba1^+^ microglia. Red: Iba1; Blue: CD68. Scale bar:10 μm. t=-7.968, P<0.001, Two-tailed unpaired t-test. **(M)** Experimental procedure to investigat the effect of overexpressing ATG7 in DG engrams and pharmacologically depleting microglia on the contextual fear memory. Mice were fed with food containing Dox (control) or food containing Dox and PLX3329 after engrams labeling. **(N)** Freezing level in context A and context B. F_treatment × context (2,45)_ = 5.406, P=0.008, RM ANOVA. Context A: EGFP versus ATG7-Flag, P<0.001; ATG7-Flag versus ATG7-Flag+PLX, P=0. 003. Bonferroni post t test. Data are presented as mean ± SEM. *P < 0.05, **P < 0.01, ***P < 0.001.

To assess the function of ATG7-dependent autophagy in DG engrams, we selectively downregulated *Atg7* in DG engrams. *AAV-RAM-Cre, AAV-DIO-Atg7-shRNA-EGFP* (or *DIO-Scramble-shRNA*) were delivered into dorsal DG to test the memory destabilization (Fig 2F). The expression of *Atg7* mRNA was dramatically decreased in *Atg7-shRNA* expressing (*Egfp^+^*) engrams (Supplemental Figure S2A-B). Downregulating ATG7 in DG engrams increased the freezing level of mice in the conditioning context (Fig 2G). No differences were detected in the open field or elevated plus maze tests, suggesting locomotor activity or anxiety level in mice were not affected by ATG7 downregulation (Supplemental Figure S2C-D). These results indicated that decreased autophagy in DG engrams may prevent memory loss.

To examine whether ATG7-dependent autophagy mediates Rac1-dependent forgetting, WT mice were infected with *AAV-RAM -Cre*, *AAV-DIO-Atg7-shRNA-EGFP* (or *scramble shRNA*), and *AAV-DIO-Pa Rac1-mVenus* (or *C450A*) bilaterally in dorsal DG (Fig 2H). Optical activation of Pa Rac1 within DG engrams led to decreased freezing level in the fear conditioned context, indicating impaired contextual fear memory, whereas downregulation of ATG7 in the DG engrams impeded the decreased freezing level (Fig 2I), suggesting ATG7 within DG engrams mediates the memory forgetting triggered by Rac1 activation.

Since ATG7 expression could be elevated after memory encoding (Fig 2E), the effects of ATG7 upregulation in DG engrams were assessed. Mice were injected with *AAV-RAM-Cre, AAV-DIO-Atg7-Flag* (or *DIO-EYFP*) in dorsal DG to specifically overexpress ATG7 in the engrams. Atg7 was specifically located in the Cre^+^ region (Fig 2J) and immunostaining of DG showed a dramatic increase of phagocytic marker CD68 puncta within Iba-1^+^ microglia in mice overexpressing ATG7 compared with the EYFP group, indicating that ATG7 in DG engrams increased the activation of the microglia (Fig 2K-L). Behavioral results showed that overexpression of ATG7 decreased the freezing level in the conditioning context (Fig 2M, N). Additionally, mice treated with diet containing Pexidartinib (PLX3397), a selective inhibitor of colony-stimulating factor 1 receptor (CSF1R) that depletes microglia (35) (Supplemental Figure S2E-F), prevented the ATG7-induced impairment of memory (Fig 2M, N). These data support the notion that autophagy and ATG7 expression in DG engrams are increased after memory encoding. Downregulation of ATG7 in DG engrams decreased the forgetting induced by Rac1 activation while overexpression of ATG7 in DG engrams increases microglia activation in DG.

### ATG7 in DG engrams induces microglia activation and immune response via TLR2/4 pathway

To investigate the mechanism of microglia activation by ATG7-dependent autophagy in DG engrams, the mice were infected with virus to specifically overexpress ATG7 in the engrams (Fig. 3A). The dorsal DG was dissected 3 days after engrams labeling, and microglia were enriched using magnetic-activated cell sorting (MACS) (Fig 3A).

**Figure 3.**
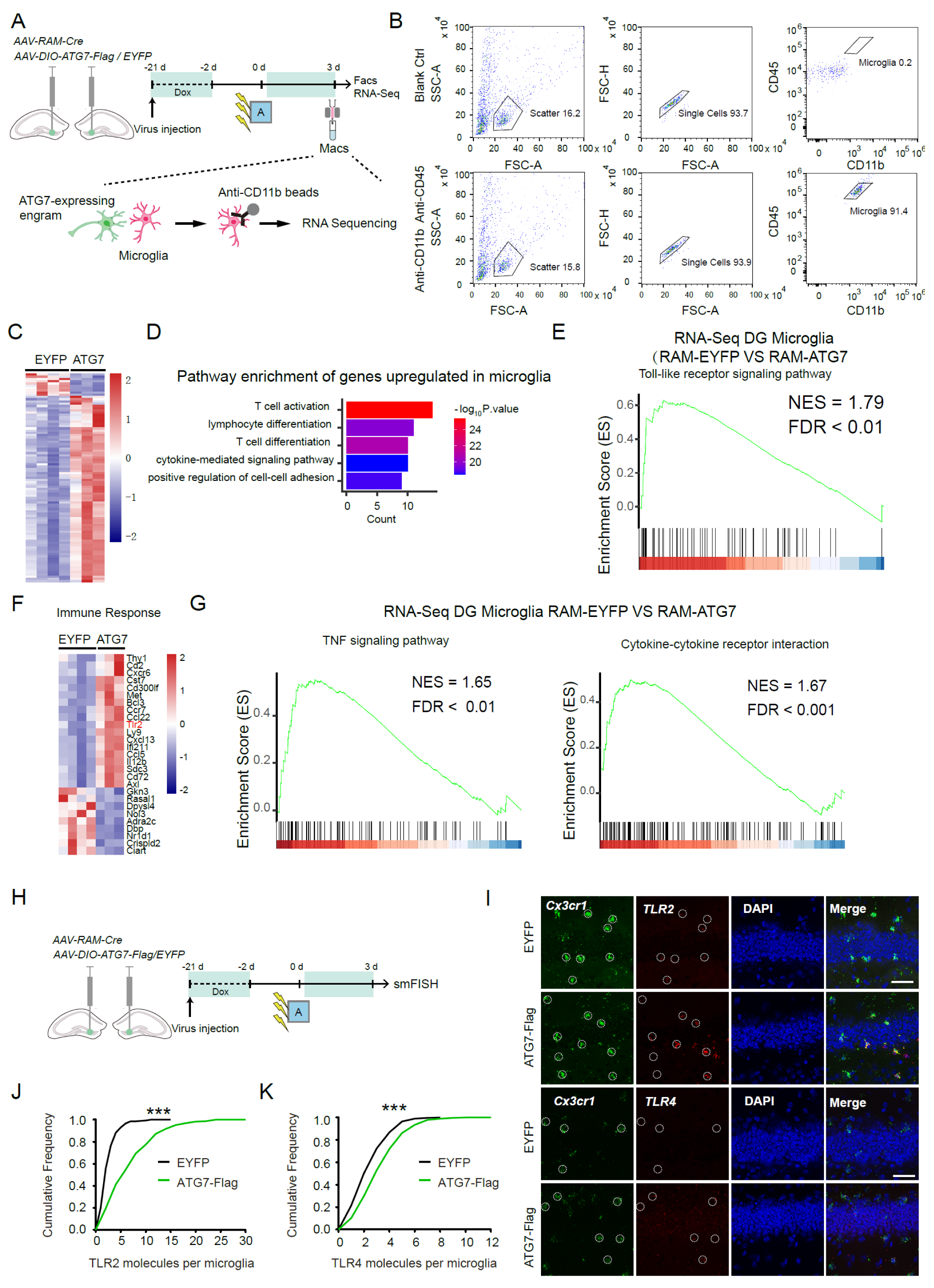
Upregulation of ATG7 in DG engrams induces immune responses and upregulates the expression of Tlr2/4 of DG microglia. **(A)** Experiment procedure. Dorsal DG bilaterally infected with AAV-*RAM-Cre* mixed with either *AAV-DIO-EGFP* or *AAV-DIO-Atg7-Flag* was dissected 3 days after fear conditioning, and the microglia were isolated by magnetic-activated cell sorting (MACs). **(B)** Fluorescence-activated cell sorting (FACS) analysis in Blank-Ctrl group (without antibody) and Anti-CD45/CD11b group (with anti-CD45/CD11b antibody) after MACs. **(C, D)** mRNA of microglia was extracted and subjected to RNA-Seq. Heat map of differentially expressed genes of the microglia **(C)** and the Gene Ontology (GO) analysis of upregulated genes in microglia **(D)** in the groups overexpressing EYFP or ATG7 within DG engrams. **(E)** GSEA results of the signature genes of theToll-like receptor (TLR) signaling pathway in the microglia when overexpressing ATG7 within DG engrams. **(F)** The heat map expression of the changed genes involved in immune response. **(G)** GSEA results of the signature genes of theTNF signaling pathway and cytokine-cytokine receptor interaction in the microglia when overexpressing ATG7 within DG engrams. **(H)** Experiment procedure to test the level of *Tlr2/4* mRNA in DG. **(I-K)** Representative images **(I)** and quantification **(J, K)** of the smFish against *Tlr2/4* mRNA in DG microglia (*Cx3cr1*^+^ cells) when overexpressing EGFP or ATG7-Flag within the DG engrams. Green: *Cx3cr1*; Red: *Tlr2/4*; Blue: DAPI. Scale bar: 50 μm. *Tlr2:* Z=5.695, P<0.001; *Tlr4*: Z=2.381, P<0.001, Kolmogorov–Smirnov test. Data are presented as mean ± SEM. \*\*\**P* < 0.001.

Fluorescence-activated cell sorting analyses were performed to assess the purity of the enriched microglia (Fig 3B). RNA-Seq and Gene Ontology (GO) of biological process categories with enrichment showed that when overexpressing ATG7 within DG engrams, the upregulated genes in DG microglia were enriched in immune response pathways, including T cell activation and cytokine-mediated signaling pathway (Fig 3C-D). Gene Set Enrichment Analysis (GSEA) revealed that the TLR signaling pathway was upregulated in microglia when engrams overexpressed ATG7, indicating the positive correlation between ATG7 overexpression within engrams and TLR signaling upregulation within microglia (Fig 3E). When analyzing the genes involved in immune response, the expression of Toll-like receptor 2 (*Tlr2*), the most abundant TLR family member expressed in microglia was increased (Fig 3F). It has been well-documented that TLR2 contributes to the activation of microglia(10, 36, 37), initiation of inflammatory signaling, and the release of Interleukin-1 alpha (IL-1α) and tumor necrosis factor alpha (TNFα), which mediates synaptic pruning(36). Consistently, GSEA showed the enrichment of genes involved TNF signal pathway and cytokine-cytokine receptor interaction in microglia when engrams overexpressed ATG7 (Fig 3G), suggesting the ATG7-dependent autophagy in the engrams promotes immune responses in microglia. To verify the expression of Tlr2 and Tlr4 in microglia, we used smFISH against *Tlr2* or *Tlr4* in *Cc3cr1* labeling microglia (Fig 3H). The expression of *Tlr2/4* mRNA in *Cx3cr1^+^* microglia of DG were upregulated by overexpression of ATG7 in the DG engrams compared with the EYFP group (Fig 3I-K), demonstrating that the TLR signaling pathway was upregulated in microglia when engrams overexpressed ATG7.

Next, we explored whether TLR2/4 within microglia were actually involved in the impaired fear memory induced by autophagy induction. *Cx3cr1-CreER* mice were infected with *AAV-RAM-Flp*, *AAV-RAM-Frt-Atg7-Flag*, and *LV-DIO-Tlr2/4miR-mCherry* (or *NegmiR-mCherry*) in dorsal DG (Fig 4A), in which Cre-loxp and Flp-FRT (38) system were combined to genetically manipulate engrams and microglia separately. After memory encoding, ATG7 was overexpressed in the engrams expressing Flp, then TAM was administered to induce the expression of Tlr2/4miR in the microglia of DG (Fig 4B). Tlr2/4 microRNA significantly downregulated the expression of *Tlr2* and *Tlr4* mRNA in DG microglia (Supplemental Figure S3A-B). The colocalization of Iba1 with mCherry indicated the specific expression of Tlr2/4miR-mCherry in DG microglia (Supplemental Figure S3C). Overexpression of ATG7 in DG engrams impaired the contextual fear memory, whereas downregulation of TLR2/4 in the DG microglia impeded the impairment of fear memory (Fig 4C).

**Figure 4.**
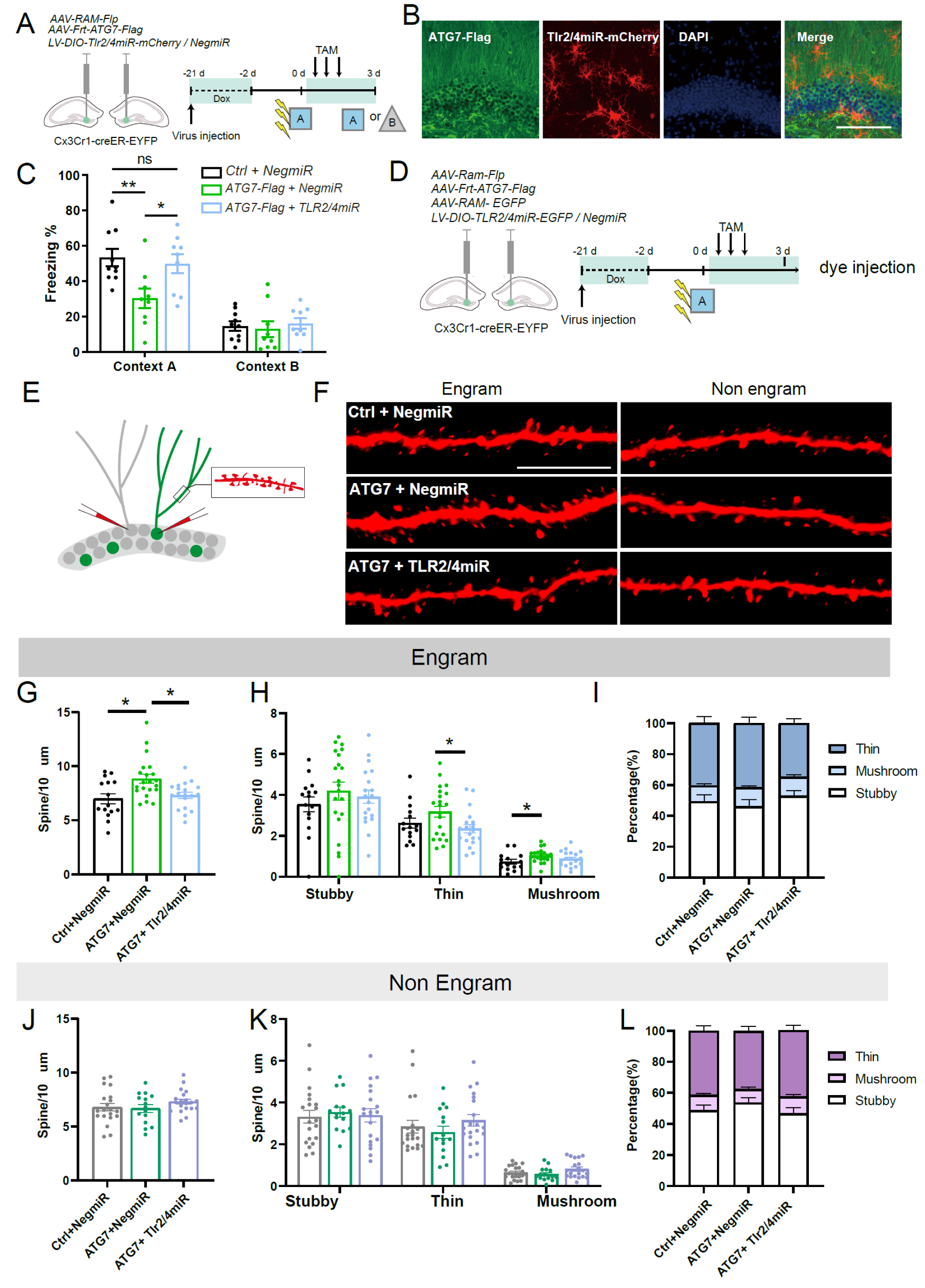
TLR2/4 in DG microglia mediates ATG7-induced spine remodeling of the DG engrams and memory impairment. **(A)** Experiment procedure to test the effect of ATG7 overexpression within DG engrams and TLR2/4 knockdown within DG microglia on the contextual fear memory. **(B)** Representative confocal images of the ATG7 and *Tlr2/4miR-mCherry* expression. Green: ATG7-Flag; Red: mCherry; Blue: DAPI. Scale bar:100 μm. **(C)** Freezing level of *Cx3cr1-CreER* mice 3 days after conditioning in context A and context B. F_treatment x context (2,25)_ = 8.517, P=0.002, RM ANOVA. Context A: EYFP+NegmiR versus ATG7-Flag + NegmiR: P<0.001; ATG7-Flag + NegmiR versus ATG7-Flag + Tlr2/4miR: P<0.001. Bonferroni post t test. **(D, E)** Experiment procedure to assess autophagy-driven synaptic elimination. **(F)** Representative confocal images of spine density within DG engrams of *Cx3cr1-CreER* mice. **(G-I)** Quantification of spine density within DG engrams of *Cx3cr1-CreER* mice. **(G)** Kruskal-Wallis H test, χ^2^=9.535, P=0.009. **(H)** Stubby: One-way ANOVA, F=0.667, P=0.517; Thin: One-way ANOVA, F=3.323, P=0.044. ATG7-Flag+NegmiR versus ATG7-Flag+Tlr2/4miR, P=0.014. Bonferroni post t test. Mushroom: One-way ANOVA, F=2.959, P=0.061. EGFP+NegmiR versus ATG7-Flag+NegmiR, P=0.019. Bonferroni post t test. **(I)** Kruskal-Wallis H test, P>0.05. **(J-L)** Quantification of spine density within DG nonengrams of *Cx3cr1-CreER* mice. **(J)** One-way ANOVA, F=1.111, P=0.337. **(K)** Stubby: One-way ANOVA, F=0.140, P=0.870; Thin: Kruskal-Wallis H test, χ^2^=3.017, P=0.221. Mushroom: Kruskal-Wallis H test, χ^2^=3.042, P=0.218. **(L)** Stubby: Kruskal-Wallis H test, χ^2^=0.701, P=0.704. Thin: One-way ANOVA, F=0.862, P=0.428. Mushroom: One-way ANOVA, F=1.000, P=0.375. Red: biocytin; Scale bar :5 μm. Data are presented as mean ± SEM. \**P* < 0.05, \*\*\**P* < 0.001.

### TLR2/4 in DG microglia mediates ATG7-induced spine remodeling of the DG engrams and memory impairment

Microglia shape brain circuits by eliminating weaker or inactive neuronal synapses that are tagged with complement proteins (3) thus underlying natural forgetting in the healthy adult brain. To assess how TLR2/4 is involved in the interaction between engrams and microglia. To investigate this question, *Cx3cr1-CreER* mice were injected with *AAV-RAM-Flp*, *AAV-RAM-Frt-Atg7-Flag*, *AAV-RAM-EGFP*, and either *LV-DIO-NegmiR-EGFP* or *LV-DIO-Tlr2/4miR-EGFP* in dorsal DG (Fig 4D). The dendrite of the engrams was labeled with EGFP. The dendritic spine morphology of engrams (EGFP^+^ neurons) and adjacent non-engrams (EGFP^-^ neurons) in DG were analyzed by single-cell microinjections with Alexa Fluor 568 hydrazide (Fig 4E-F). The total density of dendritic spines of DG engrams was increased when overexpressing ATG7, (Fig 4G). Although the proportion of each subtype of spines in DG engrams was not different (Fig 4H), the density of mushroom spines was increased after ATG7 overexpression (Fig 4I). Furthermore, downregulating the Tlr2/4 in DG microglia decreased the total spine and thin spine density in DG engrams overexpressing ATG7 (Fig 4J). In adjacent non-engrams, neither the density of total or different subtypes of spines were changed by blocking the Tlr2/4 signal in microglia (Fig 4K-L). These results suggested that ATG7 dependent autophagy in engrams might reduce spine pruning, and thus disturb established engram cell connectivity, resulting in destabilization of memory and forgetting, while downregulating TLR2/4 in microglia moderated the spine density. These results indicate that microglia promote the remodeling of the spine in a TLR2/4-dependent manner and these remodeling underlies the destabilization of fear memory.

### TLR2/4 signal in DG microglia -mediated Rac1 –dependent memory forgetting

To assess whether elevated Rac1 might activate microglia, phagocytic marker CD68 of microglia and Iba1 immunostaining were performed after optical stimulation of Rac1 (Fig 5A-B). A significantly increased intensity of CD68 puncta in the Iba1^+^ microglia was observed in Pa Rac1 group (Fig 5A-B). To investigate whether microglia activation in DG is involved in Rac1-dependent memory forgetting, we removed microglia by PLX3397 in the mice before optical activation of Rac1 to explore the effect on behavioral performance. No differences were detected in the open field or elevated plus maze tests, suggesting locomotor activity or anxiety level in mice were not affected by Rac1 activation or microglia depletion (Supplemental Figure S4A-B). 3 days after memory encoding, the decreased freezing level induced by optical activation of Pa Rac1 within DG engrams was attenuated by PLX3397 administration (Fig 5C-D).

**Figure 5.**
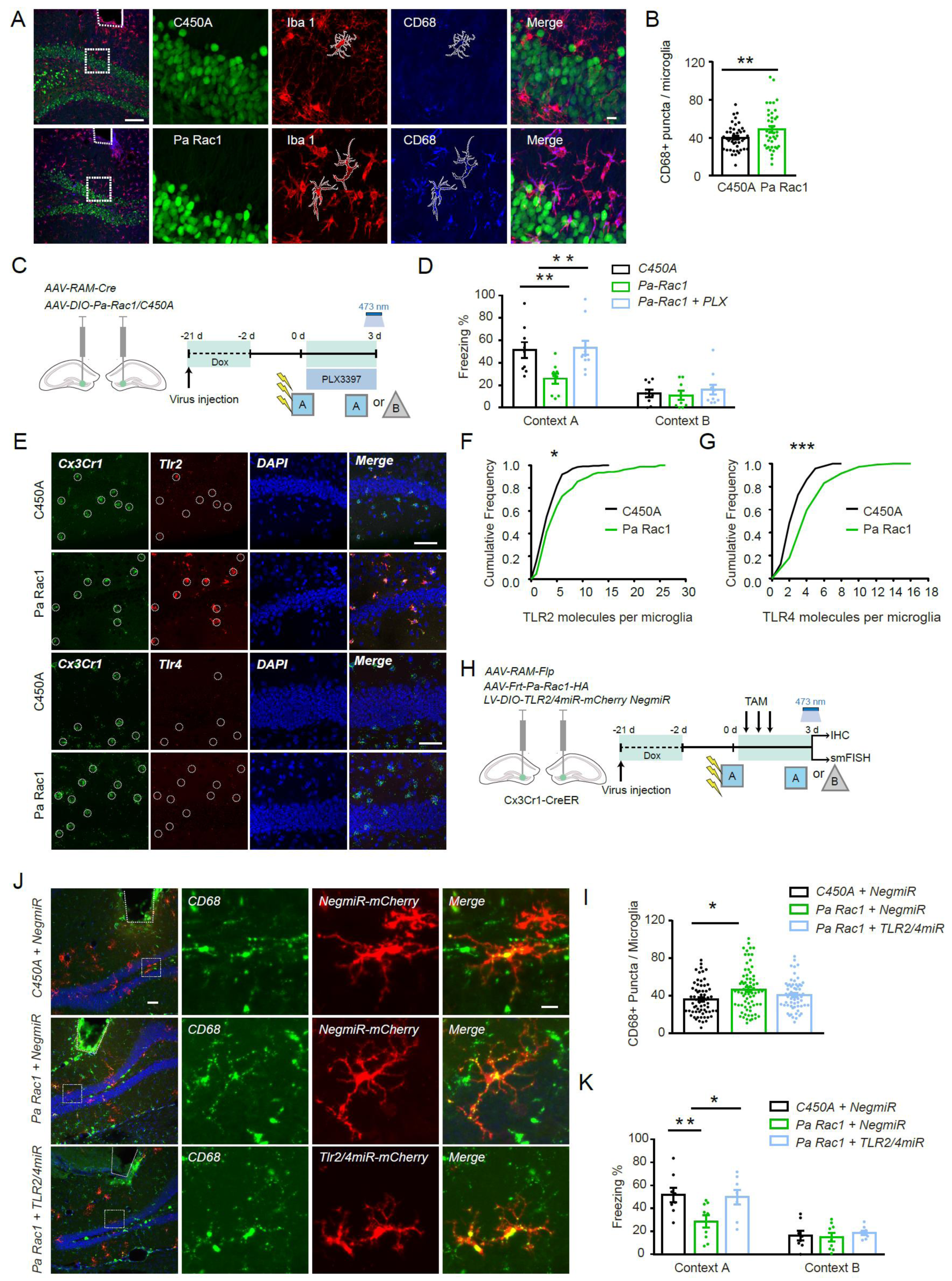
TLR2/4-dependent activation of microglia-mediated Rac1-induced memory impairment. **(A)** Representative images and quantification **(B)** of CD68^+^ puncta within microglia in C450A and Pa Rac1 group.t=-2.256, P=0.027, Two-tailed unpaired t-test. Green:C450A/Pa Rac1; Red: Iba1; Blue: CD68. Scale bar: left: 100 μm; right:10 μm. **(C)** Experiment procedure to test the effect of microglia depletion and activation of Rac1 in DG engrams on the contextual fear memory. Mice were given control food (only containing dox) or PLX food (containing dox and PLX) after engrams labeling. **(D)** Freezing level of mice 3 days after fear conditioning in context A and context B. F_treatment x context (2,26)_ = 6.403, P=0.005, RM ANOVA. Context A: C450A + Ctrl versus Pa Rac1 + Ctrl: P=0.005; Pa Rac1 + Ctrl versus Pa Rac1 + PLX: P=0.001. Bonferroni post t test. **(E, G)** Representative images **(E)** and quantification **(F-G)** of the smFish against *Tlr2/4* and *Cx3cr1^+^* mRNA in the DG. *Tlr2*: Z=1.792, P=0.003; *Tlr4*: Z=2.583, P<0.001. KS test. Green: *Cx3cr1*; Red: *Tlr2/4*; Blue: DAPI. Scale bar: top: 50 μm; bottom: 50 μm. **(H)** Experiment procedure to test the effect of Rac1 activation in DG engrams and TLR2/4 knockdown in DG microglia on the contextual fear memory. **(I)** Freezing level of *Cx3cr1-CreER* mice 3 days after fear conditioning in context A and context B. F_treatment x context (2,22)_ = 3.214, P=0.06, RM ANOVA. Context A: C450A +*NegmiR* versus Pa Rac1 + *NegmiR*: P=0.005; Pa Rac1 + *NegmiR* versus Pa Rac1 + *Tlr2/4miR*: P=0.01. Bonferroni post t test. **(J)** Representative images and quantification **(K)** of CD68+ puncta in mCherry^+^ microglia. χ2=8.184, P=0.017, Kruskal-Wallis H test. C450A +NegmiR versus Pa Rac1+NegmiR: P= 0.013. Green: *CD68*; Red: *mCherry*; Blue: DAPI. Scale bar: left:50 μm; right:10 μm. Data are presented as mean ± SEM. \**P* < 0.05, \*\**P* < 0.01, \*\*\**P* < 0.001.

smFISH following optical stimulation showed a significantly increased *Tlr2/4* mRNA level within microglia in mice overexpressing Pa Rac1 in the engrams, as compared with the C450A group (Fig 5E-G). To assess the functional interplay between Tlr2/4 in microglia and Rac1 in the engrams, *Cx3cr1-CreER* mice were infected with *AAV-RAM-Flp*, *AAV-Frt-Pa Rac1* (or *Frt-C450A*), and *LV-DIO-Tlr2/4miR-mCherry* (or *-DIO-NegmiR-mCherry*) in bilateral dorsal DG. After memory encoding, engrams were labeled with Pa Rac1. The mice were injected with TAM to downregulate TLR2/4 in microglia (38) (Fig 5H). Optical activation of Rac1 in the DG engrams significantly increased the density of CD68 puncta in the microglia surrounding the tips of the optical fibers, while knocking down TLR2/4 within microglia abolished the upregulation of CD68 puncta (Fig 5I-J). There were no differences in locomotor activity or anxiety level in mice by Rac1 activation or microglia TLR2/4 downregulation (Supplemental Figure S4C-D). However, TLR2/4 downregulation attenuated memory forgetting induced by optical activation of Rac1 within DG engrams (Fig 5K).

Collectively, these data indicate that TLR2/4-dependent microglia activation mediates Rac1-induced memory forgetting.

## Discussion

In this study, we provided a mechanistic way to elucidate the interaction between engrams and microglia that possibly contributes to memory forgetting. Specifically, we found that the expression of Rac1 in DG engrams was increased after contextual fear learning and this upregulation of Rac1 induced ATG7 expression and activated microglia, promoted spine remodeling, and thus impaired contextual fear memory. Optogenetic inhibition of Rac1 activity or downregulation of ATG7 protein both increased fear memory. Besides, TLR2/4 within microglia was essential for autophagy-dependent crosstalk between engrams and microglia. Thus, results presented here reveal a novel molecular link between the up-regulation of ATG7 in the engrams and the memory destabilization by activation of TLR pathways in the microglia.

Accumulating studies have identified mechanisms that underlie memory formation and storage benefited by the developing methods that enable more precise identification, labeling and manipulation of the neuronal components of brain circuits responsible for specific behaviors (42–44). In contrast, we know much less about the primary mechanisms governing the natural forgetting which is equally important for the memory balance. Although recent progress have begun to unravel the mechanisms underlying different forms of forgetting(45), the best characterized is Rac1-dependent forgetting which functions on the size and shape of synaptic spines by regulating actin polymerization and thus alters synaptic plasticity and behavioral memory (46, 47). Rac1 is a member of the Rho GTPase family of small G proteins which acts as key regulators of cytoskeleton dynamics and modulates lamellipodial extensions of growth cones in neurons. Studies in both Drosophila and mice have suggested that Rac1 mediates different types of active forgetting. In Drosophila, elevated Rac activity in mushroom body neurons accelerates memory decay and contributes to interference-induced forgetting. Constitutive transgenic expression of the dominant form of Rac1 in the mushroom body neurons inhibits intrinsic forgetting of odor-shock pairing(48–50). Loss-of-function mutations in autism-risk gene homologs lead to the inability to activate Rac1-dependent reversal-learning activated forgetting (51). In mice, virus-mediated expression of constitutively active Rac1 and optogenetic activation of Pa-Rac1 both cause forgetting(25, 26).

Here we showed that upregulation of Rac1 signaling in DG engram accelerated forgetting (Fig 1). Three-trials CFC paradigm increased *Rac1* mRNA level in DG engrams 12 hours after memory encoding. Moreover, different from the past to investigate the Rac1/Cofilin signaling pathway in neuron interaction, (52, 53), we found that Rac1 in engrams facilitated autophagy induction and activated microglia to destabilize memory (Fig 1 and Fig 5). This provides a novel downstream target of Rac1 signaling to regulate synaptic remodeling. However, the precise coordination of Rac1 signaling and autophagy induction is still lacking. Bernadette et al found that the TBC/Rab GAP Armus coordinates Rac1 and Rab7 functions during autophagy (54), but it remains unclear whether and how Rac1 regulates autophagy to mediate forgetting.

Autophagy is a conservative catabolic process that cleans out damaged organelles and protein aggregates engulfed in autophagosomes (APs) and degraded in lysosomes to maintain homeostasis. In the brain, autophagy is essential for the development of a healthy brain and the dysfunction of autophagy in the brain is believed to be involved in the pathogenesis of Alzheimer’s disease (AD)(37, 55, 56) and Parkinson’s disease (PD) (57, 58). As a cellular process regulating homeostasis, autophagy must adapt to the internal and external environment of the neurons, which means that the activity of autophagy itself will significantly affect the neuron state. Indeed, autophagy function in neurons to affect memory remains controversial. Emerging data suggest that autophagy in CNS plays a role in memory formation and long-term memory(34, 59–61). While there are some studies demonstrate that autophagy contributes to memory destabilization. Autophagy induction in the amygdala or in the hippocampus enhances auditory fear memory erasure via AMPAR endocytosis(20), and contributes to the degradation of the AMPAR upon LTD and GABAA receptor degradative trafficking (21, 62).

Accumulating evidence demonstrates the physiological importance of microglial autophagy in brain function and pathology. Mice lacking microglial Atg7 show various neural deficits, such as impaired synaptic pruning, enhanced Aβ-mediated inflammation(40, 68), and the correlates between autophagy and microglia activation are suggested to be involved in the pathogenesis of autism spectrum disorders and schizophrenia(69, 70). Here, we demonstrate that the autophagy in DG engrams is increased after memory encoding in a time-dependent manner (Fig 2). Downregulating ATG7, which is essential for APs formation, specifically in the engrams, increased the freezing level in the conditioning context but did not affect the locomotion activity and anxiety level (Supplemental Figure S2C-D, G-H) (63, 64).

Microglia, the predominant immune cells within the brain, have been implicated in synapse remodeling which is dependent on neuron-microglia signaling transduction. For example, cytokine interleukin-33 (IL-33) can be released from DG neurons in an expression-dependent manner and binds to the IL-33R in microglia, thus promoting microglial engulfment of the extracellular matrix and leading to the increased spine density(71). Secretion of proinflammatory cytokine IL-1β can be enhanced by induction of autophagy depending on ATG5 via an unconventional secretion pathway and then drives microglia activation(72) or induces NFκB activation(73). Toll-like receptors (TLRs) in microglia recognize the pathogen-associated molecular patterns (PAMPs) and act as endogenous ligands of the host to respond to microenvironment changes. Recent studies have shown crucial roles of TLR4-dependent activation of microglia in neurodegenerative diseases, such as AD and PD(74). Previous studies found that TLR4 activation impairs the phagocytic capacity of microglia(40) and TLR2-mediated autophagy promotes microglial cell death through microglial phenotypic switching(41).

In our study, we found that Rac1 and autophagy signals within the DG engrams both activated microglia (Fig 1 and Fig 2), providing a new mechanism for the communication between neurons and microglia. However, we didn’t examine the specific cell mediators released from DG engrams that overexpressed ATG7, but abundant researches revealed that autophagy influences the transcription and release of several cytokines(72, 75, 76) which may activate microglia. In addition, some undegraded cytoplasmic components also rely on the activation of microglia for phagocytosis. Our GSEA analysis also indicated the TNF signal pathway and cytokine-cytokine receptor interaction in microglia were upregulated in ATG7-overexpressing mice (Fig 3). Moreover, we showed that the level of TLR2/4 within DG microglia was increased and downregulating it not only refined the engram spine (Fig 4) but also rescued the impaired memory in mice overexpressing ATG7(Fig 4 and Fig 5), indicating TLR2/4 is necessary for neuron-microglia cross-talk. However, microglia are equipped with a set of other receptors, including purinergic receptors, chemokine receptors, Fc receptors, and interferon-induced transmembrane proteins, and these receptors allow microglia to recognize invading pathogens, misfolded proteins, chemokines and cytokines, metabolites, inorganic substances, and changes in pH or extracellular matrix(76). Therefore, defining the ligands and receptors between neurons and microglia is an important area for future study.

## ACKNOWLEDGEMENTS

Funding: This work was supported by grants from the Ministry of Science and Technology (2021ZD020350 to L.M. and F.W.), the Natural Science Foundation of China (31930046 & 82021002 to L.M., 31871021 to F.W.), and Shanghai Municipal Commission of Science and Technology (2018SHZDZX01 to L.M. and F.W.) and ZJLab.

## AUTHOR CONTRIBUTION

FW, LM and ZW designed the experiments and analyzed the data. WZ and RC carried out the stereotaxic surgery, behavioural tests, immunohistochemistry, and RNAscope ISH. RC and QL carried out the image quantitation. ZW constructed the viral vectors and drafted the manuscript. FW and LM revised the manuscript. FW and LM supervised the study.

## Materials and methods

### Key Resources Table

| REAGENT or RESOURCE | SOURCE | IDENTIFIER |
| --- | --- | --- |
| Antibodies |  |  |
| anti-c-Fos | Abcam | ab190289 |
| anti-Flag | Abcam | ab205606 |
| anti-CD68 | Bio-Rad | MCA1957 |
| anti-mCherry | Thermo Fisher | M11217 |
| anti-RFP | Rockland | 600-401-379 |
| anti-GFP | Rockland | 600-101-215 |
| anti-Iba1 | Wako Chemicals | 019-19741 |
| anti-p62 | Cell Signaling | 5114S |
| anti-ATG7 | Abcam | Ab53255 |
| goat anti-rabbit 488 | Jackson ImmunoResearch | 111-545-144 |
| goat anti-rabbit Cy3 | Jackson ImmunoResearch | 111-165-144 |
| donkey anti-rat 647 | Jackson ImmunoResearch | 712-605-150 |
| goat anti-rat Cy3 | Jackson ImmunoResearch | 112-165-167 |
| CD11b-PE | Thermo Fisher | 12-0112-81 |
| CD45-FITC | BioLegend | 103108 |
| Chemicals, Peptides, and Recombinant Proteins |  |  |
| PLX3397 | Selleck | S7818 |
| DAPI | Roche | 10236276001 |
| TRIzol LS | Invitrogen | 15596026 |
| Tamoxifen | Sigma | T5648 |
| doxycycline | Sigma | D3447 |
| Bacterial and Viral Strain |  |  |
| pAAV-Ef1a-DIO EYFP | addgene | 27056 |
| pTriEx-mVenus-PA-Rac1 | addgene | 22007 |
| pTriEx-mVenus-PA-Rac1-T17N | addgene | 22017 |
| pTriEx-mVenus-PA-Rac1-C450A | addgene | 22021 |
| ptfLC3 | addgene | 21074 |
| pCAG-Cre | addgene | 13775 |
| pLV-PGK-DIO-mCherry-TLR4miR-TLR2miR | addgene | 137733 |
| pLV-PGK-DIO-mCherry-nega miR | addgene | 137732 |
| pAAV-EF1a-Flpo | addgene | 55637 |
| pAAV-Ef1a-fDIO hChR2(H134R)-EYFP | addgene | 55639 |
| pAAV-RAM-EGFP | addgene | 84469 |
| pAAV-TIWB-myrmScarlet-I-P2A-post-eGRASP | addgene | 111584 |
| pAAV-CMV-bGlobin-Flex-EGFP-MIR30shRNA | Obio Technology |  |
| Critical Commercial Assays |  |  |
| RNAscope® Multiplex Fluorescent Detection Kit v2 | ACD | Cat No.323110 |
| RNAscope® 2.5 Universal Pretreatment Reagents | ACD | Cat No.322380 |
| Adult Brain Dissociation Kit | Miltenyi Biotec | 130-107-677 |
| CD11b microBeads | Miltenyi Biotec | 130-093-634 |
| PrimeScript RT reagent Kit | Takara | RR037A |
| SYBR Premix Ex Taq | Takara | RR420A |
| VAHTS mRNA-seq V3 Library Kit | Vazyme | NR611-01 |
| TLR4-C1 | ACD | Cat No. 316801 |
| ATG7-C2 | ACD | Cat No. 561261-C2 |
| EGFP-C2 | ACD | Cat No. 400281 |
| EGFP-C1 | ACD | Cat No. 400281 |
| Cre-C1 | ACD | Cat No. 402551 |
| Rac1-C1 | ACD | Cat No. 517461 |
| Cx3cr1-C2 | ACD | Cat No. 314221-C2 |
| TLR2-C1 | ACD | Cat No. 317521 |
| Slc17a6-C2 | ACD | Cat No. 319171-C2 |
| Experimental Models: Organisms/Strains |  |  |
| Cx3cr1-CreERT2 | The Jackson Laboratory | Strain: 021160 |
| Deposited data |  |  |
| Raw and processed NGS data | NCBI Gene Expression Omnibus | GSE169019 |
| Software and algorithms |  |  |
| Imaris 9.3.1 | Bitplane |  |
| SPSS 20.0 | IBM |  |

### Resource availability

Further information and requests for resources and reagents should be directed to and will be fulfilled by corresponding author.

## EXPERIMENTAL MODEL AND SUBJECT DETAILS

### Animals

*Cx3cr1-CreERT2* mice (RRID:IMSR_JAX:021160)(Parkhurst et al., 2013) were purchased from Jackson Laboratory (Sacramento, CA, USA), and were bred to C57BL/6 J for more than 6 generations. C57BL/6J male mice were obtained from the Shanghai Laboratory Animal Center (CAS, Shanghai, China) and housed in reverse light–dark cycle (lights-off at 8:00 a.m., lights-on at 8:00 p.m.) with free access to food and water. All experiment procedures for animals were performed strictly in accordance with the National Institutes of Health Guide for the Care and Use of Laboratory Animals, and were approved by the Animal Care and Use Committee of Fudan University.

## METHOD DETAILS

### Plasmid construction

The *scramble* short hairpin RNA (shRNA) (5’-TCGAAGTATTCCGCGTACGTT-3’) and shRNA targeting differentregion of mouse *Atg7* (5’-GCCAACATCCCTGGATACAAG-3’ and 5’-TTCTGTCACGGTTCGATAATG-3’) were subcloned into the *pAAV-CMV-bGlobin-DIO-EGFP-MIR30shRNA* vector (Obio Technology, Shanghai, China). Mice *Atg7* mRNA coding sequence (NM_001253717.2) tagged with FLAG at the C-terminals was synthesized by GENEWIZ (Shanghai, China), and inserted between NheI and AscI sites of *pAAV-Ef1a-DIO-EYFP* to generate *pAAV-Ef1a-DIO-Atg7-Flag*. To generate *pAAV-Ef1a-DIO-mVenus-PA-Rac1/C450A/T17N*, the EYFP in *pAAV-Ef1a-DIO-EYFP* (Addgene: No.27056) was replaced with *mVenus-PA-Rac1*, *mVenus-PA-Rac1-C450A* and m*Venus-PA-Rac1-T17N* generated from *pTriEx-mVenus-PA-Rac1* (Addgene: No. 22007), *pTriEx-mVenus-PA-Rac1-C450A* (Addgene: No. 22021) and *pTriEx-mVenus-PA-Rac1-T17N* (Addgene: No.22017), respectively. The *PA-Rac1* and *PA-Rac1-C450A* tagged with HA at the C-terminals were inserted between NheI and AscI sites of *pAAV-Ef1a-DIO-EYFP* to generate *pAAV-Ef1a-DIO-PA-Rac1/C450A-HA*. The *RFP-GFP-LC3* coding sequence was obtained from ptfLC3 (Addgene: No.21074) by PCR and inserted between NheI and AscI sites of *pAAV-Ef1a-DIO-EYFP* to replace EYFP. *PA-Rac1-HA*, *PA-Rac1-C450A-HA* and *Atg7-Flag* obtained by PCR were inserted between NheI and AscI sites of *pAAV-Ef1a-fDIO hChR*2*(H134R)-EYFP* (Addgene: No. 55639) (Fenno *et al.*, 2014a) to generate *pAAV-EF1a-fDIO-PA-Rac1/C450A-HA* and *pAAV-Ef1a-fDIO-ATG7-Flag*. The EGFP in *pAAV-RAM-EGFP* (Addgene: No. 84469) (Sorensen *et al.*, 2016) were replaced with *Flpo, Cre, or myrmScarlet-I-P2A-post-eGRASP* in *pAAV-EF1a-Flpo* (Addgene: No. 55637), *pCAG-Cre* (Addgene: No. 13775), and *pAAV-TIWB-myrmScarlet-I-P2A-post-eGRASP* (Addgene: No. 111584) (Choi *et al.*, 2018) to generate *pAAV-RAM-Flpo*, *pAAV-RAM-Cre*, and *pAAV-RAM-myrmScarlet-I-P2A-post-eGRASP*. The mCherry in *pLV-PGK-DIO-mCherry-TLR4miR-TLR2miR* (Addgene: No.137733) or *pLV-PGK-DIO-mCherry-NegmiR* (Addgene: No.137732) (Nie *et al*., 2018) were replacedwith EGFP to generate *pLV-PGK-DIO-EGFP -TLR4miR-TLR2miR* or *pLV-PGK-DIO-EGFP -Neg miR*. All AAV vectors were packaged by Obio Technology (Shanghai, China) into serotype 2/9, and the Lentiviral vectors were packaged by Genechem Co., Ltd (Shanghai, China).

### Surgery

Mice were anesthetized with 2% isoflurane during the surgery and were bilaterally injected with 500 nl of purified AAV (10^12^ IU/mL). The virus was slowly injected into DG (coordinates: ±1.4 mm mediolateral (ML), -2.0 mm anteroposterior (AP), -1.9 mm dorsoventral (DV)) with 10 μl Hamilton microsyringe (Hamilton, Reno, NV, USA) at a rate of 100 nl min^-1^. After injection, the needle was kept in place for 5 min before it was slowly withdrawn. For optical stimulation experiments, 200 μm diameter optic fiber (0.37 NA) in the cannula was bilaterally implanted in the DG with an angle of 10° from the middle to the lateral, and then were secured in place with dental cement.

### Context Fear Conditioning

Mice were habituated in the experimental room for 3 days, and the diet containing 40 mg/kg doxycycline (D3447; Sigma–Aldrich, Saint Louis, MO, USA) was taken off 48 hrs before the contextual fear conditioning. Mice were placed in the conditioning chamber illuminated with white light (Med-Associates, St. Albans, VT, USA) and housed in sound attenuating chambers. Fear conditioning was conducted in Context A (Plexiglas observation chamber with stainless-steel bars connected to a shock generator on the floor) for 300 s. The mice received three shocks (2 s duration, 0.75 mA) at 180 s, 240 s and 300 s (Guo et al., 2018). The chambers were wiped with 75% alcohol in each test interval. After the conditioning training, mice were kept back on diet containing Dox with or without PLX3397 (S7818, Selleck, Houston, TX, USA, 290 mg/kg). 3 days after training, mice were placed in the training context (context A) and a neutral context (context B, triangular chambers with white plastic floor and grey cover) for 3 min to record the freezing behavior. For optogenetic activation of Rac1 in vivo, optic fibers were connected to a 473 nm laser diode via a FC/PC adaptor (Newdoon Inc., Hangzhou, China). The laser diode was adjusted to ∼20 mW at the end of the fiber. The light pulse was delivered by a light-emitting diode (Newdoon Inc.) at 150 ms, 1 Hz for 1 hr, and the freezing behavior was measured 1 hr after stimulation. *Cx3cr1-CreER* mice were administered with Tamoxifen (T5648, Sigma–Aldrich) dissolved in ethanol and corn oil by intraperitoneal injection (150 mg/kg) for 3 consecutive days to induce Cre-dependent *TLR2/4 miR* expression in microglia.

### Elevated plus maze (EPM) test

The elevated plus maze apparatus composed of two open arms and two closed arms (34.5 cm length × 6.3 cm width × 19.5 cm height) were placed 75 cm above the floor. Mice were allowed to explore EPM freely for 6 min. Mouse behaviors were recorded with an overhead camera. The apparatuses and testing area were wiped with 75% alcohol in each test interval. Videos were analyzed with TopScan automated detection system (CleverSys. Reston, VA, USA). The percentage of time spent in the open arms and the total number of entries in the open arms were analyzed.

### Locomotor Activity Test

Mice were placed in an open field chamber (43.2 cm length × 43.2 cm width × 30.5 cm height, Med-Associates, USA) and allowed to freely explore the space for 30 min. The behavior of mice was monitored with an overhead camera. The overall activity was quantified with the TopScan automated detection system (CleverSys, Reston, VA, USA) and the total distance traveled and percentage time spent in the center zone were measured.

### Immunofluorescence and confocal microscopy

Mice were transcardially perfused with saline followed by 4% paraformaldehyde (dissolved in 1 × PBS). Brains were isolated and immersed in 4% PFA at 4 °C overnight for post fixation, and then the brains were transferred to 30% sucrose in PBS for 2 days. Brains were frozen and 40-μm coronal brain slices were prepared with a cryostat microtome (Leica CM3050S, Nussloch, Germany). For immunostaining, slices were incubated with blocking buffer (PBS containing 0.3% Triton X-100 and 10% normal donkey serum) for 1 hr at room temperature (RT) and then incubated with appropriate primary antibodies at 4 °C overnight. The primary antibody used are: anti-c-Fos (ab190289, Abcam), anti-Flag (ab205606, Abcam), anti-CD68 (MCA1957, Bio-Rad, California, USA), anti-mCherry (M11217, Thermo Fisher, Waltham, MA, USA), anti-RFP (600-401-379, Rockland, Limerick, PA, USA), anti-GFP (600-101-215, Rockland), anti-Iba1 (019-19741, Wako Chemicals, VA, USA), anti-ATG7 (ab53255, Abcam). After rinsed in PBS at RT, the slices were incubated with secondary antibodies for 1 hr and then stained with DAPI (10236276001, Roche, Basel-Stadt, Schweiz) for 10 min at RT. The secondary antibodies used are from Jackson ImmunoResearch (West Grove, PA, USA): goat anti-rabbit 488 (111-545-144), goat anti-rabbit Cy3 (111-165-144), donkey anti-rat 647 (712-605-150), goat anti-rat Cy3 (112-165-167). Slices were extensively washed in PBS and mounted onto glass slides in Fluor Mount-G mounting medium (0100-01, Southern Biotech, Birmingham, AL, USA).

Fluorescent images were captured by Nikon-A1 confocal laser scanning microscope (Tokyo, Japan) with ×20 or ×60 objectives for imaging. The center of viral infection was taken as the brightest fluorescent point. The tip of the fiber was determined by the ∼50-μm-thick gliosis generated by the fiber. The slides for smFISH were photographed using a Nikon A1 confocal laser scanning microscope and a ×20 objective lens with 3 × optical zoom. For analyzing CD68 puncta and immunofluorescent intensity, images of the microglia within 200 μm from the optical fiber tips were acquired with a 20 × objective lens with 5 × optical zoom. All images were analyzed with open-source software Image J.

### Microglia isolation, Fluorescence-activated cell sorting analysis and RNA sequencing

The mice were perfused with ice-cold 1× PBS and hippocampi were isolated and incubated with enzyme mix (130-107-677, Miltenyi Biotec, Bergisch Gladbach, Germany) to dissociate into single-cell suspension. Debris was removed using debris removal solution (130-107-677, Miltenyi Biotec). Microglia were enriched using CD11b microBeads (130-093-634, Miltenyi Biotec). For fluorescence-activated cell sorting analysis, cells were stained with CD11b-PE (12-0112-81, Thermo Fisher) and CD45-FITC antibodies (103108, BioLegend, San Diego, CA, USA) and sorted on CytoFLEX S Flow Cytometer (Beckman Coulter, Brea, CA, USA). For RNA sequencing, RNA in microglia was extracted with Trizol (15596026, Thermo Fisher). Libraries were prepared following the instructions in VAHTS mRNA-seq V3 Library Kit (NR611-01, Vazyme, Nanjing, China) for illumina.

### RNA sequencing analysis

The quality of reads was evaluated using Fastp (Jiang et al., 2021), all samples passed quality control, and reads were aligned to mm10 (GRCm38 from Ensembl) using Hisat2 (Kim et al., 2019). Mapped reads were counted using featureCounts (Wang et al., 2020b) and the DESeq2 package (Niu et al., 2018) was used to perform differential gene expression analysis. Gene ontology term enrichment analysis was performed using clusterProfiler (Yu et al., 2012). Gene set enrichment analysis of KEGG was carried out by gseKEGG (part of clusterProfiler) and the results of GSEA were visualized with gseaplot2 (created by Guangchuang Yu). The data have been submitted to NCBI Gene Expression Omnibus (GEO) under accession number GSE169019.

### Quantitative RT-PCR

To downregulate Tlr2/4 in microglia, *Cx3cr1-CreER* mice were bilaterally injected with *LV-DIO-NegmiR-mCherry* or *LV-DIO-Tlr2/4miR -mCherry* in the dorsal dentate gyrus. Two weeks later, the mice were intraperitoneally injected with tamoxifen (T5648, Sigma-Aldrich) at a dose of 150 mg/kg for 3 consecutive days. Dorsal DG was isolated and the microglia were purified using magnetic-activated cell sorting and mRNA in microglia was extracted with Trizol. cDNA was obtained from total RNA using PrimeScript RT reagent Kit (RR037A, Takara, Shiga, Japan) and then subjected to quantitative RT-PCR with SYBR Premix Ex Taq (RR420A, Takara). The primers used for PCR are as follows: *Gapdh forward*, 5’-GTGGAGTCATACTGGAACATGTAG-3’; *reverse*, 5’-AATGGTGAAGGTCGGTGTG-3’; *Tlr2 forward*, 5’-TGG AATGTCACCAGGCTGC-3’; *reverse* 5’-GTCCGTGGAAATGGTGGC-3; *Tlr4 forward*: 5’-ATGGAAAAGCCT CGA ATCCT-3’; *reverse*, 5’-TCCAAGTTGCCGTTTCTTGT-3’.

### Single-molecule RNA FISH

PFA-fixed brains were frozen in OCT and then sliced in 10 μm thick coronal sections mounted on a Colorfrost Plus slides (Thermo Scientific) followed by air-dry at room temperature (RT) overnight. Slices were performed target-retrieval and proteolysis using RNAscope® 2.5 Universal Pretreatment Reagents (ACD: 322380) In brief, the sections were incubated with hydrogen peroxide for 10 min at RT and proteolysis for 30 min at 40 °C. SmFISH probes for genes examined: Tlr2 -C1 (ACD: 317521), Tlr4-C1 (ACD: 316801), Cx3cr1-C2 (ACD: 314221-C2), Rac1-C1 (ACD: 517461), ATG7-C2 (ACD: 561261-C2), EGFP-C2 (ACD:400281), EGFP-C1 (ACD:400281), Cre-C1 (ACD:402551), Slc17a6-C2 (ACD: 319171-C2). The slices were hybridized at 40 °C for 2 hrs. Subsequent amplify signal steps were performed using the RNAscope Multiplex Fluorescent Reagent Kit v2 (ACD: 323110).

### Western Blotting

Mice were anesthetized and perfused at different time points (1hour, Day1, Day3, Day7, Day16) following context fear conditioning in context A to investigate the protein level of ATG7 or p62. Dorsal hippocampi were dissected within 5min on ice. Tissue samples were lysed for 10min in RIPA lysis buffer (10 mM tris-HCl, pH 7.4, 1% Triton X-100, 50 mM NaCl, 1.0 mM EGTA, 1.0 mM EDTA, 50 mM NaF, 100 μM Na3VO4, 1.0 μM phenylmethylsulfonyl fluoride, 1.0 mM dithiothreitol, and protease inhibitors from Roche cocktail tablets) and centrifuged at 12000g for 30min at 4 ℃. Samples were diluted in SDS-polyacrylamide gel electrophoresis loading buffer (0.3 mM tris-HCl, pH 6.8, 30% glycerol, 10% SDS, 6% β-mercaptoethanol, and 0.012% w/v bromophenol blue) and heated for 5 min at 95 ℃. Samples were loaded on 10 % SDS polyacrylamide gradient gel and transferred onto the nitrocellulose membrane. The blots were blocked with Tris-buffered saline with Tween (TBST)-5% BSA for an hour at room temperature, incubated with either anti-β-actin (A2066, Sigma) or anti-p62 (5114S, Cell Signaling) overnight at 4 ℃ and the secondary antibody for 1 hour at room temperature. The membrane was scanned with appropriate channels (Odyssey, LI-COR Biosciences) and selected films were quantified with open-source software Image J. β-actin bands were used for normalization and a control group was used as a comparison.

### Spine Morphology analysis

Mice were anesthetized and perfused with 1% PFA for 1min followed by 4% PFA for 15min. The Brains were isolated and immersed in 4% PFA at 4 °C overnight for post fixation . After post-fixation, the brain was sliced into 200-μm thickness containing DG with a vibratome (HM760V, Thermo) and immersed in 0.1M phosphate buffer at 4℃. The cell body in DG was filled with Alexa Fluor 568 hydrazide for 10 min of diffusion by using a micromanipulator and then the brain slices were mounted on slides. For analyzing dendritic spines, images were acquired by a 60 × oil objective lens with 5 × optical zoom. Z stacks were acquired at 0.5 μm intervals. Reconstruction and analysis of the dendrite synaptic structures were carried out by Imaris software (Bitplane, St. Paul, MN, USA). Apical dendritic segments of dye-filled DG cells were chosen. 50- to 80- μm segments beginning >50 μm and ending <150 μm distal to the soma and after the first branch point were quantified. Spine types were classified in accordance with the following criteria: thin (length< 2μm, mean width [head] ≧ mean width [neck] ); mushroom (length< 2μm and maximum width [head] ≧2 × mean width [neck]); stubby(length< 1μm); filopodia( length≧2μm ).To determine the spine density, 15-25 neurons from 3 mice were measured and the spine density of each type was expressed as the number of spines per 10 μm of dendrite.

### Statistical analysis

Data were analyzed with SPSS 22 software (IBM, Armonk, NY, USA), and plotted by Graphpad Prism. Our sample sizes were based on our previous research (Jiang *et al.*, 2021; Niu *et al.*, 2018; Wang *et al.*, 2020b). The normality test and homogeneity of variance test were performed by the Shapiro-Wilk test and Levene’s test. Tukey’s multiple comparisons test was performed after the student’s t test (Unpaired, two-tailed), or one-factor analysis of variance (ANOVA) followed. Bonferroni’s *post hoc* analysis was performed after repeated-measures (RM) ANOVA. Data that does not fit a normal distribution were analyzed with a nonparametric test. Two-sample *Kolmogorov Smirnov* test was used for cumulative frequency plot analysis. Statistical significance was represented as *P<0.05; **P<0.01, and ***P<0.001. All data are presented as mean ± SEM.

---

**Figure S1.**
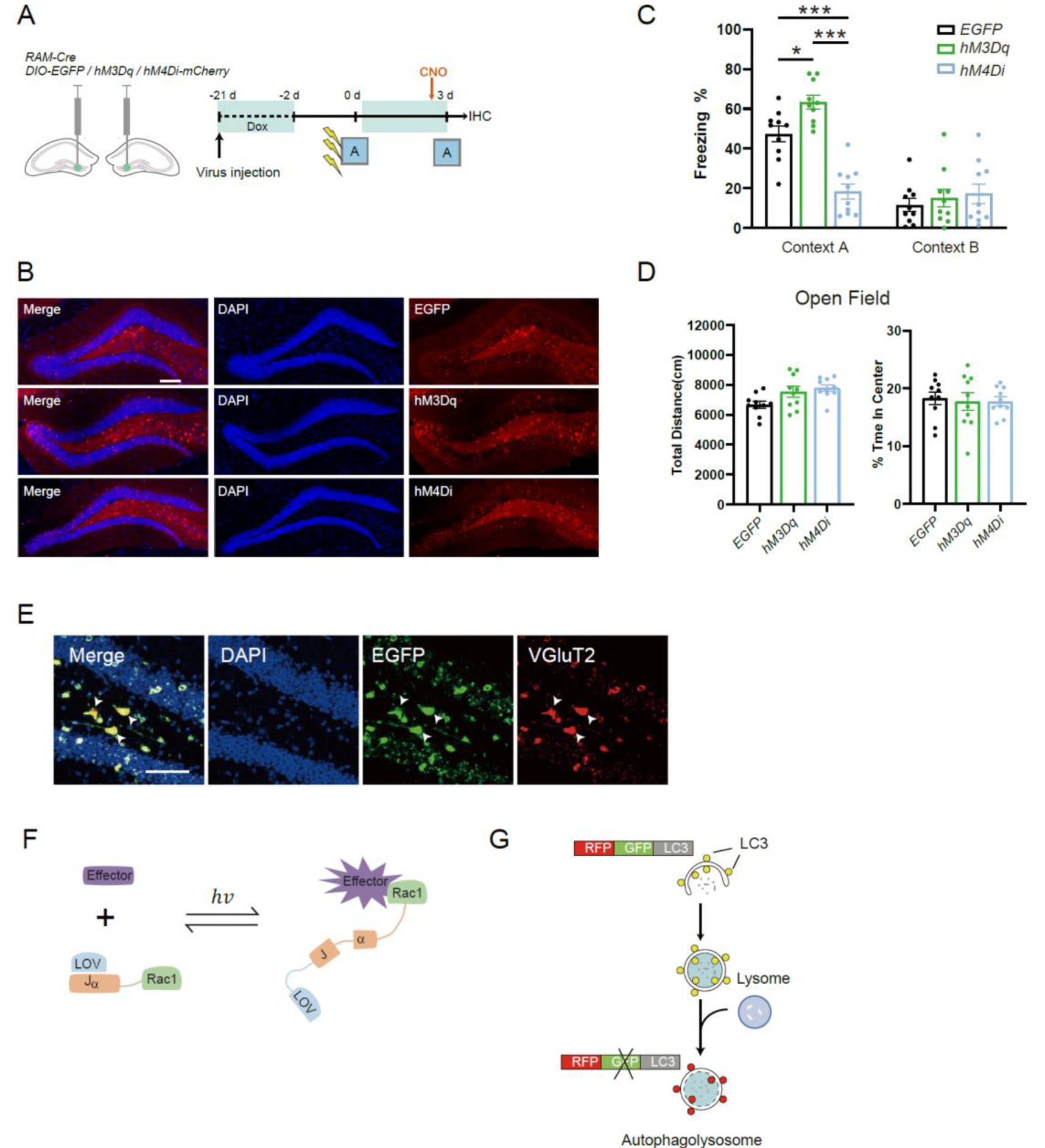
The verification of RAM system by chemogenetics and the schematic of optical-activated Rac1 system. **(A)** Experiment procedure to verified the robust activity marking (RAM) system. Clozapine N-oxide (CNO) was delivered by intraperitoneal (i.p.) injection (10mg/kg) 30 min before memory test. **(B)** Representative confocal image showing expression of chemogenetic tools labeled ensemble. Red: EGFP or mCherry; Blue: DAPI. Scale bar:200 μm. **(C)** Freezing level of mice 3 days after fear conditioning in context A and context B. F_treatment x context (2, 27)_ =22.18, P<0.001, RM ANOVA. Context A: Ctrl *vs* hM3Dq: P=0.017; Ctrl *vs* hM4Di: P<0.001. Bonferroni post t test. **(D)** The effect of engramss activation or inhibition on locomotor activity. One-way ANOVA. P>0.05. **(E)** Representative confocal images of vGluT2 expression in *Egfp*+ engrams. Green: EYFP; Red: vGluT2; Blue: DAPI. Scale bar:50 μm. **(F)** Schematic of optical-activated Rac1 system. **(G)** Schematic of Cre-dependent autophagy flux reporter system *AAV-DIO-RFP-GFP-LC3*. Data are presented as mean ± SEM. \**P* < 0.05, \*\**P* < 0.01, \*\*\**P* < 0.001.

**Figure S2.**
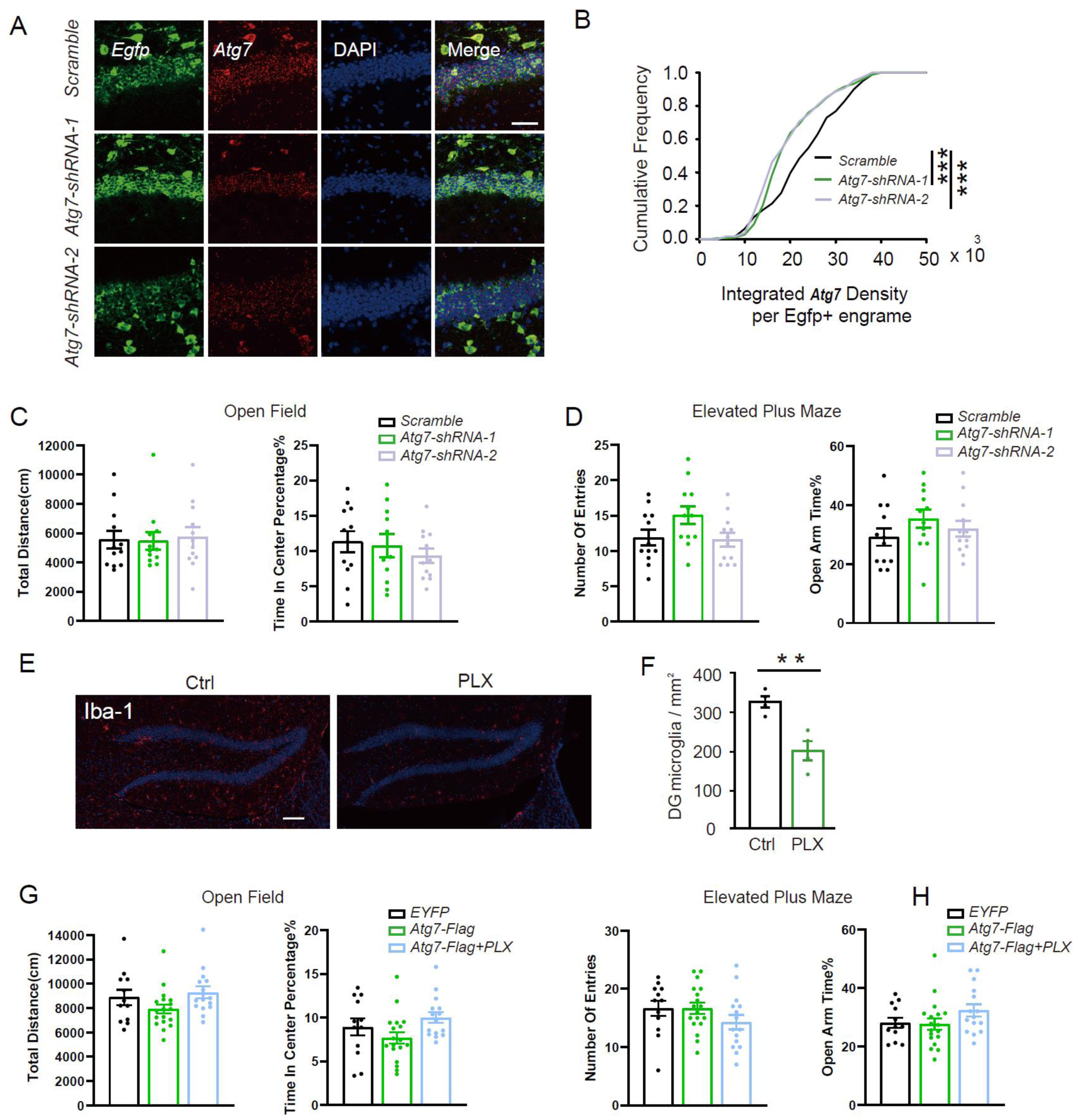
The intervention effect of *Atg7-shRNA* and Pexidartinib (PLX3397). **(A, B)** Representative confocal images **(A)** and quantification **(B)** of smFish against *Atg7* and *Egfp* mRNA to assess the knockdown efficiency of Atg7-shRNA within *Egfp*+ engrams. Green: *Egfp*; Red: *Atg7*; Blue: DAPI. Scale bar: 50 μm. **(C, D)** The effect of ATG7 downregulation in DG engrams on the Open Field **(C)** and Elevated Plus Maze (EPM) tests **(D). (E, F)** Representative confocal images **(E)** and quantification **(F)** of Iba1^+^ microglia number in the DG. Red: Iba1; Blue: DAPI. Scale bar:100 μm. t=4.308, P=0.005, Two-tailed unpaired t-test. **(G, H)** The effect of ATG7 overexpression in the DG on locomotor activity **(G)** and anxiety **(H).** Data are presented as mean ± SEM. \**P* < 0.05, \*\**P* < 0.01, \*\*\**P* < 0.001.

**Figure S3.**
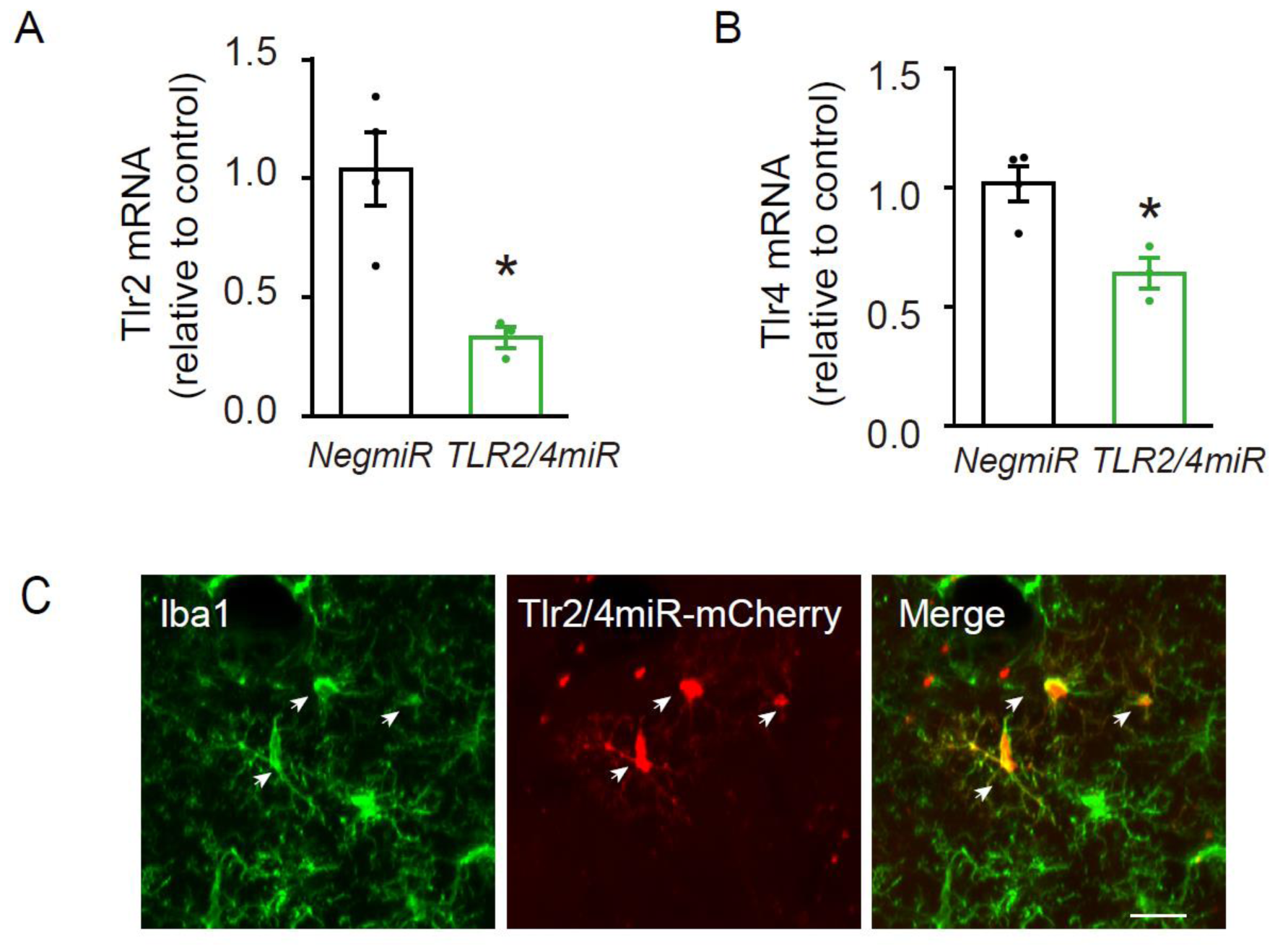
The verification knockdown efficiency of *Tlr2/4 microRNA* (A, B) *Cx3cr1-CreER* mice infected with *LV-DIO-NegmiR-mCherry* or *LV-DIO-Tlr2/4miR -mCherry* in dorsal DG was administered with TAM 1 day after engrams labeling to downregulate Tlr2/4 in DG microglia. Microglia in the DG were isolated using MACs, and the mRNA was extracted and subjected to qRT-PCR to quantify the mRNA level of *Tlr2/4* in the microglia. Tlr2: t=3.794, P=0.013; Tlr4: t=3.631, P=0.015. Two-tailed unpaired t-test. **(C)** Representative images of lentivirus expression in DG microglia. Green: Iba1; Red: mCherry. Scale bar:20 μm. Data are presented as mean ± SEM. *P < 0.05, **P < 0.01, ***P< 0.001.

**Figure S4.**
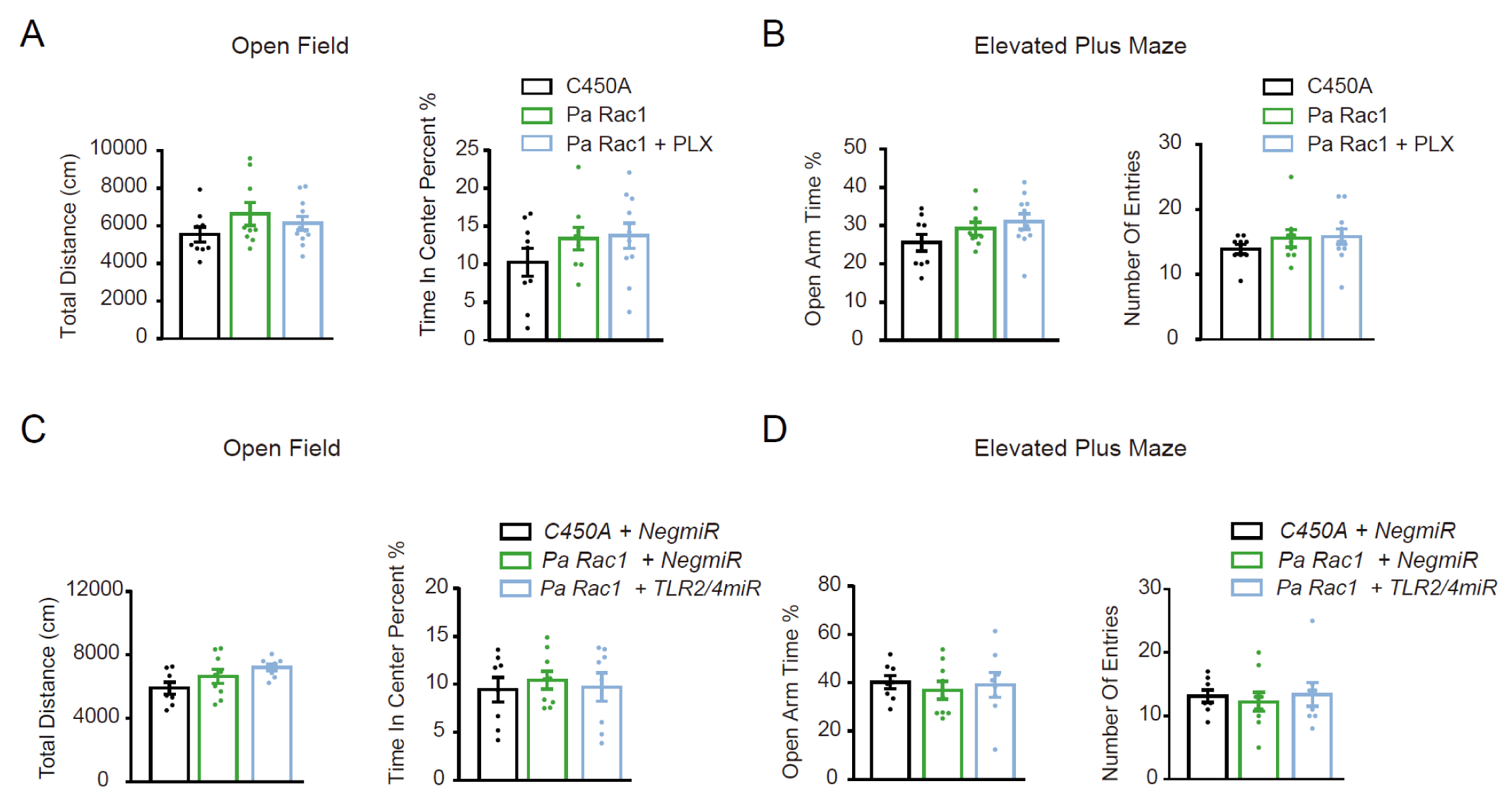
Optical activation of Rac1 and microglia manipulation in DG did not affect the locomotor activity and anxiety level of mice. **(A, B)** The effect of Rac1 activation in the DG engrams and microglia depletion on the Open Field **(A)** and EPM tests **(B)** of *Cx3cr1-CreER* mice. One-way ANOVA and Kruskal-Wallis H test, P>0.05. **(C-D)** The effect of optical-activating Rac1 in DG engrams and knocking down Tlr2/4 of DG microglia on Open Field **(C)** and EPM tests **(D)** of *Cx3cr1-CreER* mice. One-way ANOVA, P>0.05. Data are presented as mean ± SEM. *P < 0.05, **P < 0.01, ***P< 0.001.

